# Musicians’ brains at rest: Multilayer network analysis of MEG data

**DOI:** 10.1101/2024.01.02.573886

**Authors:** Kanad N Mandke, Prejaas Tewarie, Peyman Adjamian, Martin Schürmann, Jil Meier

**Author notes:** J.M. and M.S. contributed equally to this work. Corresponding author: Dr Martin Schürmann, School of Psychology, University of Nottingham, University Park, Nottingham NG7 2RD, UK. E-Mail-addresses of all other authors: Kanad Manke, Prejaas Tewarie, Peyman Adjamian, Jil Meier.

## Abstract

The ability to proficiently play a musical instrument requires a fine-grained synchronisation between several sensorimotor and cognitive brain regions. Previous studies have demonstrated that the brain undergoes functional changes with musical training, identifiable also in resting-state data. These studies analysed fMRI or electrophysiological frequency-specific brain networks in isolation. While the analysis of such “mono-layer” networks has proven useful, it fails to capture the complexities of multiple interacting networks. To this end, we applied a multilayer network framework for analysing publicly available data (Open MEG Archive) obtained with magnetoencephalography (MEG). We investigated resting-state differences between participants with musical training (n=31) and those without (n=31). While single-layer analysis did not demonstrate any group differences, multilayer analysis revealed that musicians show a modular organisation that spans visuomotor and frontotemporal areas, known to be involved in musical performance execution, which is significantly different from non-musicians. Differences between the two groups are primarily observed in the theta (6.5-8Hz), alpha1 (8.5-10Hz) and beta1 (12.5-16Hz) frequency bands.

We demonstrate that the multilayer method provides additional information that single-layer analysis cannot. Overall, the multilayer network method provides a unique opportunity to explore the pan-spectral nature of oscillatory networks, with studies of brain plasticity as a potential future application.

## Introduction

Musical proficiency is a commonly used model to study brain plasticity (Herholz & Zatorre, 2012; Jäncke, 2009b; Zatorre & McGill, 2005). This research is largely motivated by the fact that achieving high levels of proficiency in any given musical instrument requires years of focused training. Numerous studies have reported structural brain changes due to musical training (Amunts et al., 1997; Elbert et al., 1995; Schlaug et al., 1995). For example, an increased size of the anterior half of the corpus callosum (Schlaug et al., 1995), and differences in the cortical representation of the hand motor area (Amunts et al., 1997; Elbert et al., 1995). More specifically, the brain also undergoes anatomical plasticity that pertains to the demands posed by the particular type of musical training, i.e., pianists showed a left hemisphere dominance, whereas violinists showed right hemisphere advantage (Bangert & Schlaug, 2006). Robust changes in grey matter specific to auditory, motor and visuo-spatial regions have also been reported (Gaser & Schlaug, 2003). This anatomical specialisation is potentially the result of years of musical training that requires fine-grained communication between several cortical areas. For example, musicians showed an increased coupling between auditory and pre-motor areas in functional magnetic resonance imaging (fMRI) (Grahn & Rowe, 2009), related to musicians’ skilled actions during a performance.

It is worth noting that the training-related plastic changes are not unique to musicians but can be seen in groups of subjects with some degree of behavioural specialisation. For instance, training-related white matter changes have been observed in healthy adults learning to juggle (Scholz et al., 2010), professional golfers (Jäncke, 2009a), and London taxi drivers (Maguire et al., 2000).

To better characterise relationships between anatomical regions from task-free data, resting-state functional connectivity approaches have proven useful. It must be noted that either a seed-based or whole-brain functional connectivity analysis can inform about the strengths of these connections (Friston, 2011). Going a step further, studying the pattern of these connections can give us more information about the functional network as a whole. Using a seed-based connectivity approach to resting-state fMRI in musicians, demonstrated an increased functional connectivity in auditory, visual and motor areas. Similarly, (Tanaka & Kirino, 2017) demonstrated an increased functional connectivity in the supplementary motor area (SMA) network in musicians imagining musical performances based on fMRI data. Using a whole-brain analysis of fMRI, (Alluri et al., 2017) reported that musicians recruit action-based brain networks, while non-musicians recruit perception-based networks in a naturalistic listening paradigm. In a seed-based fMRI study by (Alluri et al., 2015a), musicians demonstrated an increased connectivity with SMA and with ventromedial and ventrolateral cerebral and cerebellar affective regions while listening to music, whereas non-musicians displayed a higher connectivity only with subcortical regions. In a resting-state fMRI study, (Luo et al., 2014) demonstrated a greater local functional connectivity in musicians compared to non-musicians in the bilateral dorsal anterior cingulate cortex, anterior insula, and anterior temporoparietal junction. In a similar study, increased insular connectivity in musicians was reported (Zamorano et al., 2017). As reviewed above, evidence of functional specialisation in musicians comes from structural data (white or grey matter changes) or haemodynamic functional data. Little attention has been paid so far to the analysis of electrophysiological magnetoencephalography (MEG) data recorded from musicians’ brains during resting state, capturing spontaneous, non-task-related brain activity, which could offer insights into the brain activity of musicians and the communication between brain areas on shorter temporal time scales. Activity recorded by MEG predominantly consists of neuronal oscillations, which occur over a wide range of temporal scales (1-200Hz). Numerous studies have shown that these oscillations are involved in mediating both long-range and local communication in the brain (e.g., (Fries, 2005)).

Analysis of resting-state networks is not limited only to functional magnetic resonance imaging (fMRI) but has also been a significant part of the human electrophysiological research based on MEG (Brookes et al., 2011; Hillebrand et al., 2012) and electroencephalography (EEG) (Smit et al., 2008; Stam et al., 2005). However, to the best of our knowledge, only one study (Klein et al., 2016) has explored resting-state differences between musicians and non-musicians using an electrophysiological approach. The authors analysed high-density EEG resting-state data obtained from musicians with graph-theoretical approaches. They reported that musicians showed an increased intra- and inter-hemispheric functional connectivity between auditory, sensorimotor, and prefrontal cortex in theta (6.5-8Hz), alpha1 (8.5-10Hz), and alpha2 (10.5-12Hz) frequency ranges. A limitation of this study is that separate frequency specific networks were studied in isolation (also referred to as analysis of single layers), which ignores the pan-spectral picture of ongoing brain connectivity.

To overcome this limitation, the present study used a multilayer network framework, combining data obtained from multiple frequency bands in one single network description (Brookes et al., 2016a; Mandke et al., 2018; Tewarie, Hillebrand, et al., 2016). Generally, a multilayer network is a network of networks, which is made up of individual networks as layers that are interconnected via some relationship (a more formal introduction to multilayer networks follows in the methods section). Multilayer network approaches have been introduced in the field of neuroscience where different layers of the network correspond to different M/EEG frequency bands (frequency specific networks) or networks obtained from different neuroimaging modalities (Bassett et al., 2010; Brookes et al., 2016b; Buldú & Porter, 2018; De Domenico, 2017; Mandke et al., 2018; Tewarie, Hillebrand, et al., 2016; Vaiana & Muldoon, 2020; Yu et al., 2017). Using frequency-based multilayer networks, (De Domenico et al., 2016) showed that multilayer networks can better classify between schizophrenic patients and healthy controls than single-layer or aggregated single-layer networks. Other disease-oriented applications of multilayer networks are MEG-based frequency-specific multilayer networks (Yu et al., 2017) and a multimodal diffusion weighted imaging (DWI) -fMRI-MEG multilayer network (Guillon et al., 2017) to investigate Alzheimer’s disease, where multilayer network metrics of patients in the latter study predicted individual cognitive and memory impairment.

Most complex networks – including multilayer networks – exhibit a certain level of modularity, i.e. densely connected groups of nodes (brain regions) form clusters (modules). Numerous neuroimaging studies have reliably shown that human brain architecture is highly modular (Power et al., 2011; Yeo et al., 2011). Recently, (Puxeddu et al., 2021) tested different community detection algorithms on multilayer EEG networks. Modules have been suggested to sub-serve information processing (Sporns & Betzel, 2016). The modular organisation of the brain begins early in life and matures during the adolescent years (Baum et al., 2017). This development of modular organisation allows for cortical specialisation and that reorganisation occurs with learning (Bassett et al., 2010). Accordingly, as a result of training, brain networks in musicians may reorganise to facilitate communication between sensory, motor and cognitive areas of the brain.

In the present methods-focused study, we investigated resting-state MEG recordings from musicians and non-musicians obtained from the Open MEG Archive (Niso et al., 2016). The primary aim of this study was to identify neuronal signatures of behavioural specialisation or plasticity-induced changes following musical training in MEG resting-state networks using multilayer network analysis. Furthermore, we also investigated changes in functional connectivity in motor regions that have previously shown structural and functional changes (in task-based data) following musical training (Alluri et al., 2015b; Tanaka & Kirino, 2017). In the present study, we estimated functional connectivity using amplitude envelope correlations between anatomically pre-defined regions. In a previous publication (Mandke et al., 2018), we demonstrated that group comparisons using multilayer networks can be biased by differences in link weights (e.g., by differences in average connectivity). This was ameliorated by normalising the link weights using the singular value decomposition (SVD).

We used a seed-based connectivity analysis, an established single-layer analysis method (Network-Based Statistics toolbox, (Zalesky et al., 2010)), and a multilayer network approach, with layers representing MEG frequency bands. The absence of a statistically significant difference between groups in single-layer analysis and a presence of a difference in multilayer analysis would highlight the added value of the latter approach. Here, we focused on identifying the modularity of resting-state networks within and between individual MEG frequency bands ("pan-spectral" modularity). We hypothesised that the years of musical training would have changed the modular structure of the musicians in comparison with non-musicians, where we expected sensory, motor, and cognitive areas of the brain to be stronger interconnected in musicians than in non-musicians, leading to less modularity in musicians than in non-musicians. Finally, we conclude with a discussion of some of the methodological issues in the analysis of multilayer networks.

## Methods

### Participants and MEG data pre-processing

All MEG data analysed in this study were obtained from a public access data repository, Open MEG Archive, OMEGA (Niso et al., 2016). At the time the analysis was performed, the OMEGA repository contained MEG data from 150 healthy controls. The repository also includes self-report questionnaire data. Out of the questionnaire data, we used self-reported musical experience to classify the healthy participants into two groups. We used the 5-min resting-state MEG recordings acquired on a 275-channel CTF-MEG system (MISL, Coquitlam, Canada) as detailed in (Niso et al., 2016). The first group included data from participants who had no history of neuropsychiatric disorders and 5+ years of experience playing some musical instrument (Supplementary Table 1). A total of 31 participants met this inclusion criteria. Out of the 31 participants, 23 answered “Yes” to the questionnaire item, “Do you consider yourself to be a musician?” and 8 answered “No”. For simplicity, we will refer to the whole group of 31 participants as musicians to reflect their 5+ years of experience. This group was a heterogeneous sample, as it included a wide range of musical expertise playing instruments (e.g. string instruments, percussions etc.). The second (control) group included data from neurologically healthy OMEGA participants. We identified 31 (age- and gender-matched) participants for our control group (non-musicians). Further data from the total 62 participants (31 in each group, all right-handed by self-report) are given in Table 1.

**Table 1.** Summary of statistics for both groups.

|  | *Musicians* | *Non-musicians* |
| --- | --- | --- |
| *Gender* | M: 20; F: 11 | M: 20; F: 11 |
| *Age* | Mean: 28.25 years;<br>SD: 7.25 | Mean: 27.48 years;<br>SD: 5.95 |
| *Years of experience* | Mean: 13.25 years;<br>SD: 5.04 | no experience |

*M: male; F: female; SD: standard deviation.*

The raw data included task-free, 5-mins eyes open resting-state MEG scans, sampled at 2400Hz (Niso et al., 2016). As part of our pre-processing, the data were down-sampled at 600Hz. A third order synthetic gradiometer configuration of the CTF-MEG system was applied; a 150Hz low pass anti-aliasing filter was used. OMEGA repository also provides participant’s digitised head shape (Polhemus). Using in-house software, we performed surface matching of the digitised head shape to an equivalent head shape extracted from individual anatomical scans to allow for co-registration of MEG sensor geometry and brain anatomy.

### Source localisation using an atlas based beamformer

An atlas based beamforming approach (Hillebrand et al., 2012) – previously also used in (Brookes et al., 2016b; Tewarie, Bright, et al., 2016; Tewarie, Hillebrand, et al., 2016) – was applied for source localisation. The cortex was parcellated using the automated anatomical labelling (AAL) (Tzourio-Mazoyer et al., 2002). Out of the 116 AAL regions, we selected 78 cortical regions that provide a full cortical coverage and ignored the remaining 38 subcortical and cerebellar regions in line with previous multilayer network studies (Brookes et al., 2016b; Tewarie, Bright, et al., 2016; Tewarie, Hillebrand, et al., 2016). A beamformer approach (Robinson & Vrba, 1999) was employed to generate a single time course of electrophysiological activity within each of these regions. To achieve this, the centre of mass was first derived for each region. Given the spatial resolution of MEG (Barnes et al., 2004), voxels were defined on a regular 4 mm grid covering the entire region, and the beamformer-estimated time-course of electrical activity was derived for each voxel. To generate a single time course representing the whole region, individual voxel signals were weighted according to their distance from the centre of mass using a Gaussian weighting function. This procedure ensures that the regional time course is biased towards the centre of the region, with a full width half maximum of approximately 17mm. To calculate individual voxel time courses, a scalar beamformer was used (Robinson & Vrba, 1999). Covariance was computed within a 0.5-150 Hz frequency window and a time window spanning the entire experiment (300s) (Brookes et al., 2008). Regularisation was applied to the data covariance matrix using the Tikhonov method with a regularisation parameter equal to 5% of the maximum eigenvalue of the unregularised covariance matrix. The forward model was based upon a dipole approximation (Sarvas, 1987) and a multiple local sphere head model (Huang et al., 1999). Dipole orientation was determined using a non-linear search for optimum signal to noise ratio (SNR). Beamformer time courses were sign flipped where necessary to account for the arbitrary polarity introduced by the beamformer source orientation estimation. This process resulted in 78 electrophysiological time courses, each representative of a separate AAL region. This approach was applied to each subject individually.

### Functional connectivity calculation

The time courses were frequency filtered in theta (6.5-8HzHz), alpha1 (8.5-10Hz), alpha2 (10.5-12Hz), beta1 (12.5-16Hz), beta2 (16.5-20Hz), this frequency selection was motivated by the work on training related plasticity (Klein et al., 2016; Langer et al., 2013). After frequency filtering, to correct for signal leakage, frequency filtered time courses were subjected to a multivariate orthogonalisation (Colclough et al., 2015). Using the Hilbert transform, an amplitude envelope was computed by calculating the absolute value of the analytical signal. The envelopes of all the frequency bands were down-sampled to 4Hz (Brookes et al., 2011; Luckhoo et al., 2012). Finally, a functional connectivity matrix was reconstructed for individual frequency bands by computing Pearson’s correlation coefficient between down-sampled envelope pairs, with each correlation coefficient forming a single element in the weighted adjacency matrix (O’Neill et al., 2015; Tewarie, Hillebrand, et al., 2016; van Wijk, 2014). This produced a square (78 nodes x 78 nodes) off-diagonal weighted adjacency matrix for each frequency band under consideration.

### Seed-based connectivity

Furthermore, to explore seed-based connectivity, the following regions were used – left and right SMA (nodes 12 and 51 in AAL atlas) and left and right precentral gyrus (nodes 14 and 53 in AAL atlas) (review: (Jäncke, 2009b). A linearly constrained minimum variance beamformer was used to project data to these regions. Following this, amplitude envelope correlations were used to estimate functional connectivity between these regions across participants.

### Single layer analysis using Network-based statistics

To test if the effects of musical training can be detected in raw adjacency matrices, we performed single layer analysis using the Network-Based Statistics (NBS toolbox, (Zalesky et al., 2010). For technical details on NBS, we would like to redirect the reader to the original article (Zalesky et al., 2010). A short summary of NBS is as follows: In a typical cross-sectional design, NBS identifies significantly different pairwise associations between groups, where such associations are links or connections between two nodes. A given symmetric *N x N* connectivity matrix has maximum *N(N-1)/2* individual connections. Using *M* independent permutations, a t-statistic is estimated for all individual connections. Then links that exceed a certain threshold with their respective *t* statistic are retained, which forms the supra-threshold network. The size of the biggest connected component found in this resulting network is used as a test statistic. The *p*-value of an observed component is calculated by finding the number of permutations for which the maximal component size exceeds the original test statistic, normalised by *M*. To identify group differences using NBS, we used M=5000 permutations with independent samples t-test (*p*<0.05) and tested a range of t-statistic thresholds t=0.5:0.1:5.0.

### Multilayer network construction

A multilayer network is a complex network structure, which is a network of networks, where each layer is formed by an individual network. The links in such a network describe relationships between combinations of all possible nodes and layers. In this study, multilayer networks were constructed by integrating information from theta (6.5-8Hz), alpha1 (8.5-10Hz), alpha2 (10.5-12Hz), beta1 (12.5-16Hz), beta2 (16.5-20Hz) frequency bands, where each network or layer shared the same number of nodes (78 cortical regions). The links in each layer were the amplitude envelope correlation (AEC) weights within that frequency band. Figure 1 shows a schematic of multilayer network construction.

**Figure 1.**
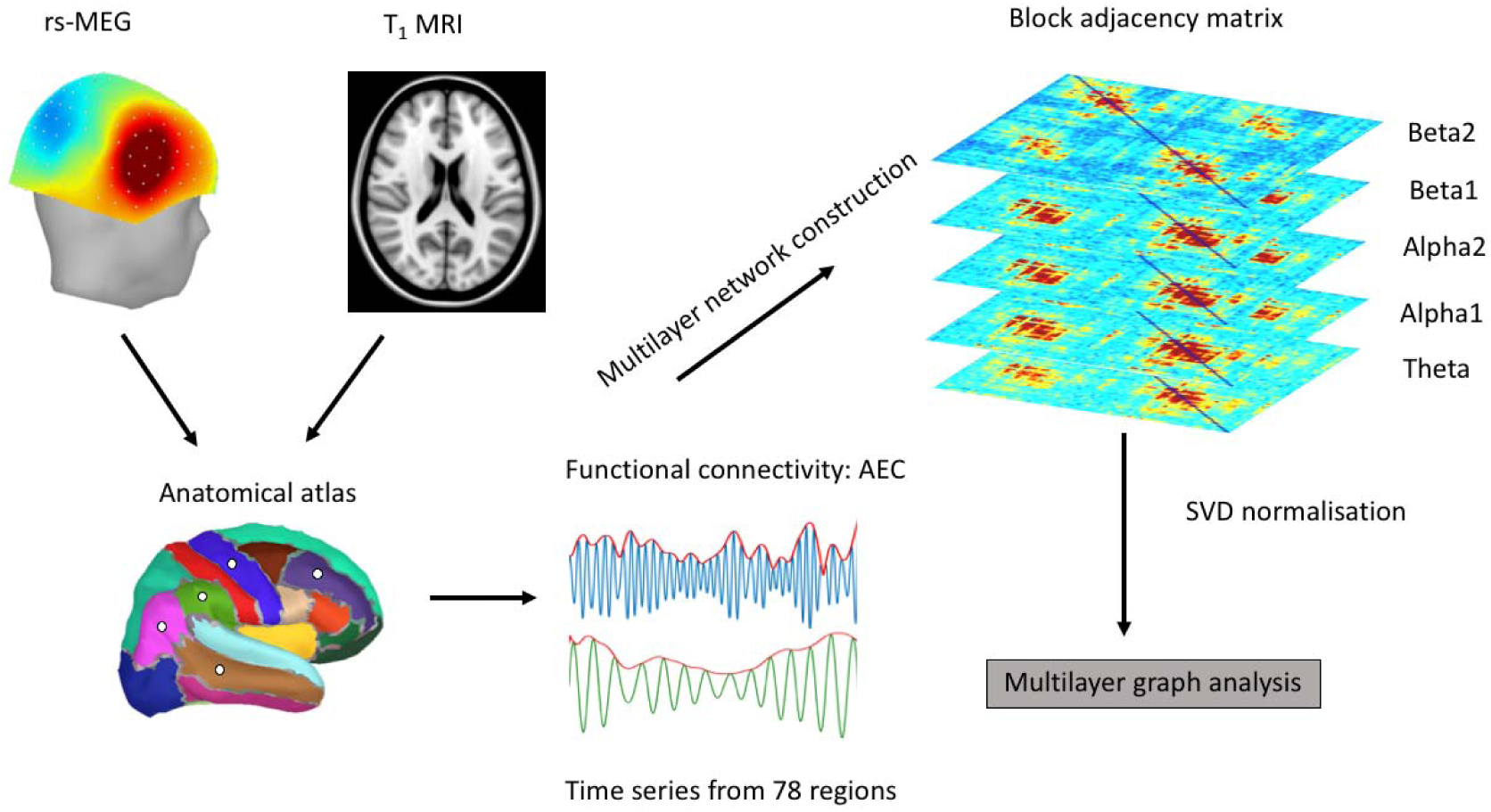
A schematic of the analysis pipeline used in the construction of multilayer networks. Here, we use a special case of the multilayer networks: multilayer networks with one-to-one between-layer coupling. rs: Resting state; AEC: Amplitude envelope correlations; Theta: 6.5-8Hz; Alpha1: 8.5-10Hz; Alpha2: 10.5-12Hz; Beta1: 12.5-16Hz; Beta2: 16.5-20Hz.

Mathematical representation of a multilayer network is its block adjacency matrix (De Domenico et al., 2013; Van Mieghem, 2016). An *f*-layered multilayer network can be written in the form of a block adjacency matrix (Sahneh et al., 2015) as follows:

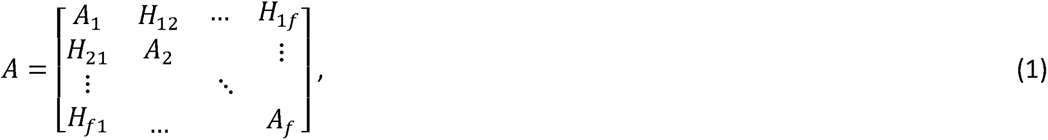

where *A_α_* corresponds to a symmetric, square adjacency matrix of a layer *α,* 1 ≤ *α* ≤ *f*. In our case *A_α_* corresponds to individual frequency specific networks. The matrix *H_kl_* corresponds to the coupling matrix between the layers *k* and *l*, where 1 ≤ *k,l* ≤ *f,f* = 5. The frequency specific networks, *A_α_*, have the same dimensions for all layers (*N x N, N=78*), implying that every layer of the network has the same number of nodes or brain regions.

To be in line with previous studies (Mandke et al., 2018; Yu et al., 2017), we chose one-to-one between layer relationship *H_kl_* = *cl*, where c is a constant and *I* the identity matrix (more details on the inter-layer coupling in Supplementary material 1). Thus, we used a case where coupling matrices are special diagonal matrices (*H_kl_* = *cl*). In other words, we allow only for links between the same nodes across all layers (i.e. we ignore cross-frequency coupling between non-identical areas in different layers). We computed all results over a range of values for the interlayer coupling, c=0:0.01:1.

### Graph analysis of multilayer networks

A multilayer network was constructed for every participant in the two groups. The absolute value was taken of all link weights. However, comparing raw block-adjacency matrices between groups can be biased due to differences in link weights, i.e. such a comparison can inflate false positives or negatives (Mandke et al., 2018). These biases in the context of multilayer networks arise due to differences in average connectivity, which can influence the calculated network metrics (van den Heuvel et al., 2017). To correct for such biases we used singular value decomposition (SVD) based normalisation approach (Mandke et al., 2018). Prior to subjecting the block adjacency matrix to graph analysis, all singular values were normalised by the largest singular value of the weighted adjacency matrix to correct for difference in mean connectivity between groups. For a given block adjacency matrix *A*, we apply a singular value decomposition *A* = *UΛV^T^*, where *U* and *V* contain the left and right singular vectors and *Λ* the singular values of *A*. To correct for differences in average connectivity, we rescaled *Λ* by the largest singular value 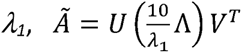 (the justification of the choice of eigenvalue can be found in the Supplementary Material 2). The rescaled matrix would thus become *Ã*. The multiplication by 10 is used to ensure that the range of values in *Ã* is not too small and varies between 0 and 1.

To identify community structure specific to musicians we used a community detection algorithm. Community detection algorithms (e.g. (Blondel et al., 2008) have widely been used to study resting-state fMRI networks and structural data (Crossley et al., 2013; Garcia et al., 2018; Sporns & Betzel, 2016). Generally, communities in a network are obtained by optimising a quality function, e.g. modularity (Newman, 2006), which provides an index for within-community and between-community connections. The multilayer modularity as expressed in (Bassett et al., 2013) is as follows:

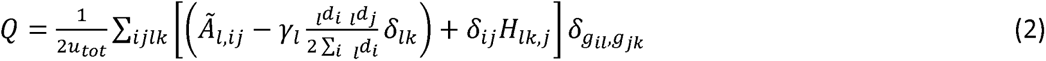

where *u_tot_* is the total link weight in the network, *γ_l_* is the resolution parameter (set to 1, constant over all subjects), *g_il_* is the community assignment of node *i* in layer *l*, and *δ_ij_* is the Kronecker delta function (meaning that *δ_ij_*=1 if *i=j* and *δ_ij_*=0 if i≠j). Using this community detection algorithm, we calculated individual nodal module assignments expressed as indices. Nodes with the same index can be said to be part of the same community. To ensure that the community organization was stable and not based on initial settings, we repeated the algorithm 100 times at every instance.

We first performed this community detection algorithm on the multilayer matrix of each individual subject. Using these index assignments, we calculated an agreement (consensus) matrix (*N x N*) for every layer per group, where each element in the matrix corresponds to the proportional number of times two given nodes were part of the same community over all individual subjects (Baum et al., 2017). Furthermore, we obtained a collapsed agreement matrix by averaging over all individual layer agreement matrices. This step was performed to achieve a group-level representation of the entire multilayer network.

To compare the multilayer community structure between groups (musicians vs non-musicians), we applied the community detection algorithm 100 times (Equation 2) to each collapsed agreement matrix, generating 100 partitions of the 78 nodes into various communities (Baum et al., 2017). We then repeated the process of constructing an agreement matrix from these 100 partitions and reapplying the community detection algorithm until achieving a stable community assignment across all 100 Louvain algorithm repetitions. This process yielded a single community assignment for each interlayer coupling value c. Next, we calculated the frequency with which each node was assigned to each module across all c values.

For some of the analyses, we used the Brain Connectivity Toolbox (https://sites.google.com/site/bctnet/, (Rubinov & Sporns, 2010)). All our used code is publicly available at https://github.com/jilmeier/multilayer-musicians/.

## Results

### Functional connectivity and Network-Based Statistics

Seed-based connectivity analysis for the 4 nodes under consideration (bilateral SMA and pre-central gyri) did not reveal any statistically significant (*p<0.05*) differences between the two groups.

After performing SVD normalisation, for visual representation of the data, functional connectivity estimates between all nodes for all frequency bands (theta, alpha1, alpha2, beta1, beta2) obtained using AEC were averaged across participants (Figure 2: musicians and Figure 3: non-musicians). Only the top 40% (i.e., the strongest) of connections were subsequently plotted in brain space to visualise the underlying networks (see Figures 2 and 3). After computing individual connectivity matrices, we performed single layer analysis using NBS. However, this analysis failed to show any significant effect with stringent statistical thresholding (*p<0.05*, FDR-corrected) across frequency bands for a large range of t-statistic thresholds. A null result using NBS is possible because of several reasons: 1) Single layer analysis ignores the pan-spectral picture in the data, which warrants a more integrative approach. 2) High differences in link weights can smear underlying experimental effects and this demands some corrections as highlighted in (Mandke et al., 2018; van Wijk et al., 2010).

**Figure 2.**
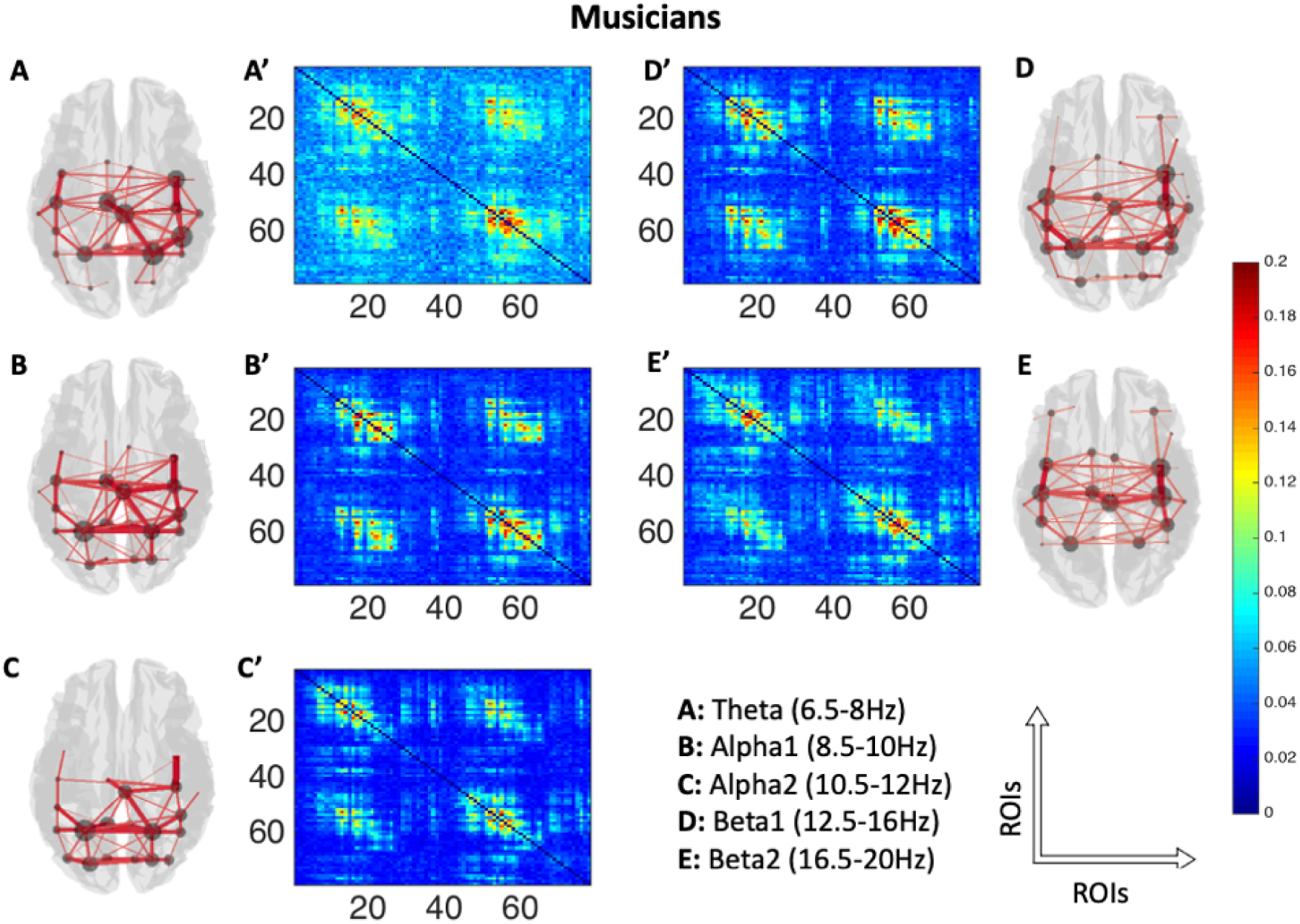
Averaged AEC in adjacency matrix (78 brain area labels on each axis: 1-39 left hemisphere, 40-78 right hemisphere) across musicians (A’ to E’) and its corresponding source space (top 40% (i.e., the strongest) of connections) representation (3D plots, A to E) in the 5 frequency bands under consideration. All adjacency matrices show typical patterns of high-AEC within left hemisphere (upper left quadrant), within right hemisphere (lower right quadrant) and between hemispheres (upper right quadrant). In the 3D plots, red lines indicate presence of connections, with thicker lines corresponding to stronger connections. The grey circles in the 3D plots indicate summed magnitude of link weights (node strength, weighted degree) between the node and the rest of the brain. ROIs: Regions of interest.

**Figure 3.**
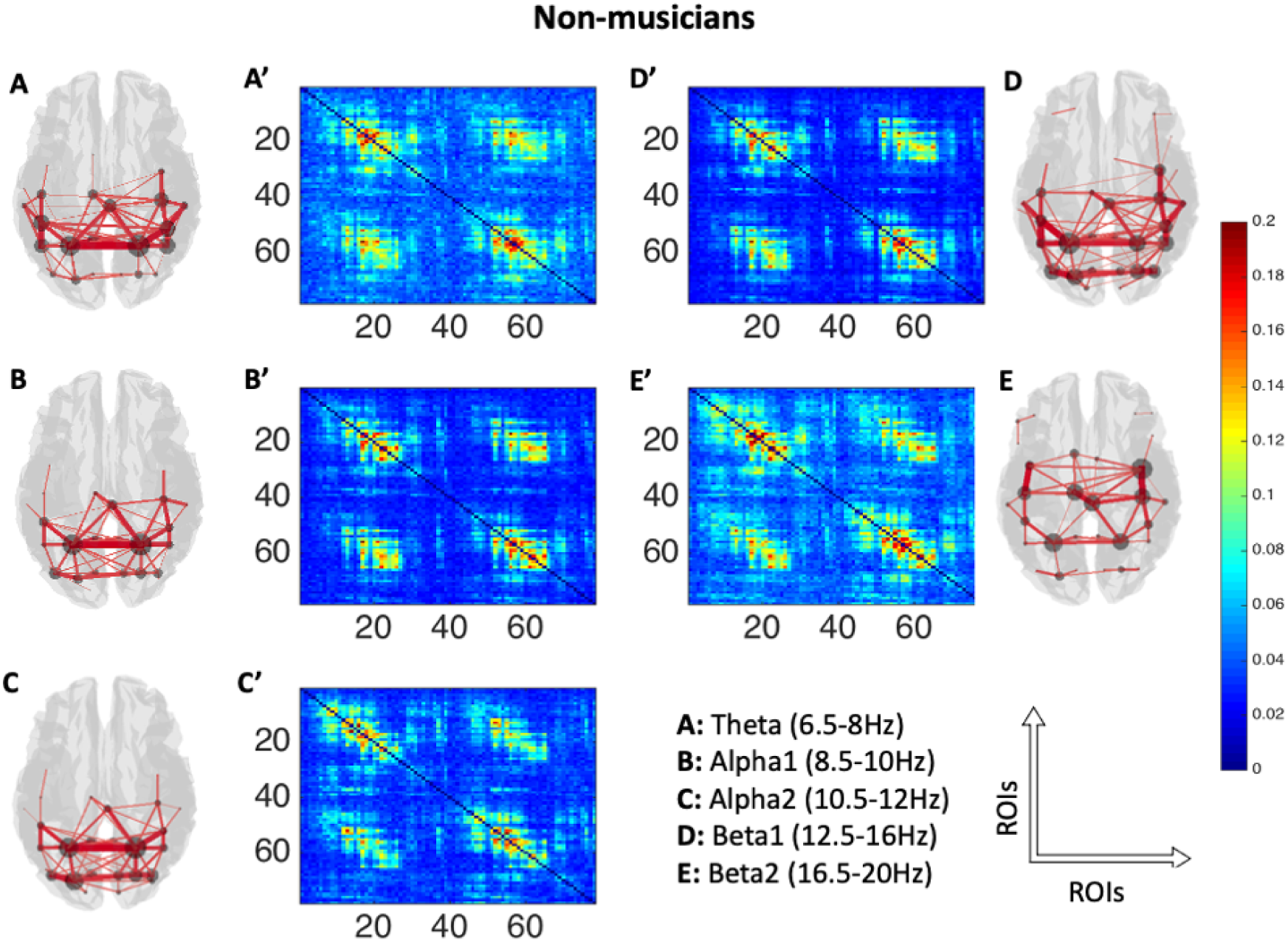
Averaged AEC plotted as adjacency matrices (78 brain area labels on each axis: 1-39 left hemisphere, 40-78 right hemisphere) across non-musicians (A’ to E’) and their corresponding source space (top 40% (i.e., the strongest) of connections) representation (3D plots, A to E) in 5 frequency bands under consideration. In the 3D plots, red lines indicate presence of connections, with thicker lines corresponding to stronger connections. The grey circles in the 3D plots indicate summed magnitude of link weights (node strength, weighted degree) between the node and rest of the brain. ROIs: Regions of interest.

#### Graph analysis of the multilayer networks: Community structure

The block adjacency matrices of the two groups were subjected to community detection algorithm after SVD normalisation. Modularity (Q) was calculated for individual subjects’ SVD normalised multilayer networks (Figure 4) and was significantly different between groups over a large range of interlayer coupling values (p<0.001 for each value of c=0:0.01:1, Wilcoxon rank-sum test). The Wilcoxon rank-sum test was conducted to compare modularity between musicians and non-musicians. The analysis yielded statistically significant results for all comparisons, with z-values ranging from −1.70 to - 6.76, p<0.001. The mean effect size for the tests, calculated as rank-biserial correlation (r), was r=−0.553, indicating a strong negative effect. Cohen’s d equivalent for the mean effect size was −1.33, further confirming a very large effect size. The negative sign indicates that the first group in the comparisons (i.e., musicians) always had lower values. A predictive classifier based on multilayer network feature of modularity confirmed that we can predict whether an individual is a musician or not with above chance level (above 54%), which increased up to 100% for increasing values of c (Supplementary Figure S1). If we consider the average modularity over all c values per individual as a classifying feature, the accuracy for prediction was 96.78% (93.44% in a 4-fold cross-validation, more details in Supplementary Material 3).

**Figure 4:**
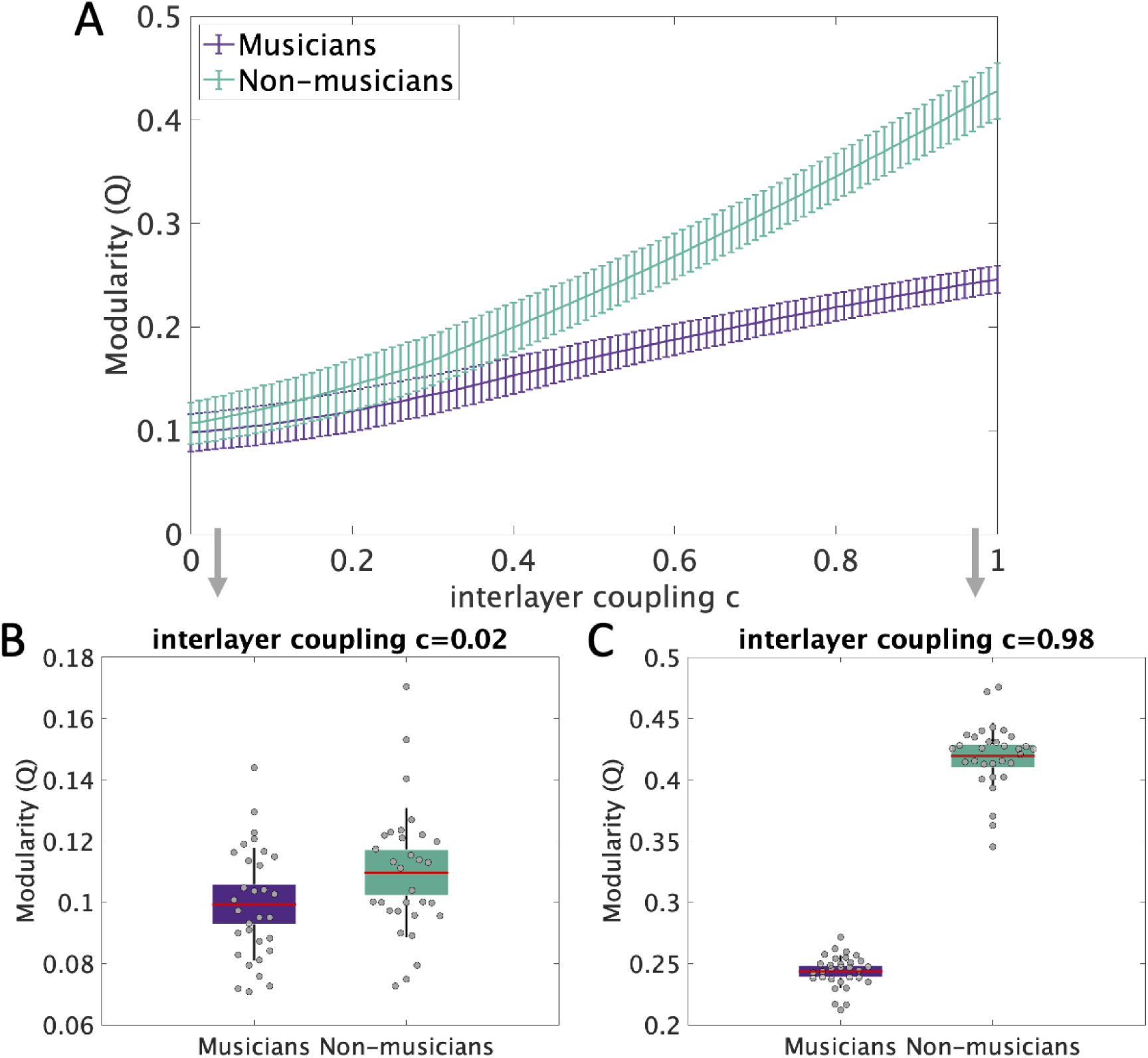
Modularity difference between musicians and non-musicians. **(A)** Modularity (Q) in the two groups of musicians (purple) and non-musicians (green), depicted as mean values with error bars in the length of their standard deviation over all 100 runs of the community detection algorithm and for each c in the range of 0:0.01:1. This difference was found to be statistically significant for all values of c (p<0.001 Wilcoxon rank-sum test). **(B-C)** Example boxplots of modularity (Q) for the two groups for c=0.02 (B) and c=0.98 (C). The boxes show 95% SEM and SD plotted as whiskers.

The group of musicians demonstrated different modular organisation than the non-musicians (average adjusted Rand index over all c values: 0.6876). The group representation of this modular organisation for both groups is shown in Figure 5. The multilayer network community structure observed in musicians had two large communities, while non-musicians had three communities. Musicians showed a modular structure that encompasses both visual and motor networks, shown in yellow (Figure 5, left panel), and frontal and temporal regions, shown in purple. In non-musicians, the modularity analysis revealed three modules, the frontal (colour: purple), midline-temporal (colour: green) and posterior (colour: yellow) network components. The non-musicians showed distinctly different community structure from the group of musicians. In non-musicians, the visual and motor networks are segregated from one another (as shown in yellow and green, respectively, see Figure 5 right panel), whereas musicians display integrated visuo-motor networks (as shown in yellow in left panel in Figure 5). In addition, the musicians also exhibit markedly different patterns of connectivity spanning the fronto-temporal regions, in particular in the right hemisphere, with integration of frontal and superior temporal regions into a single community only in musicians (Figure 5, left panel).

**Figure 5:**
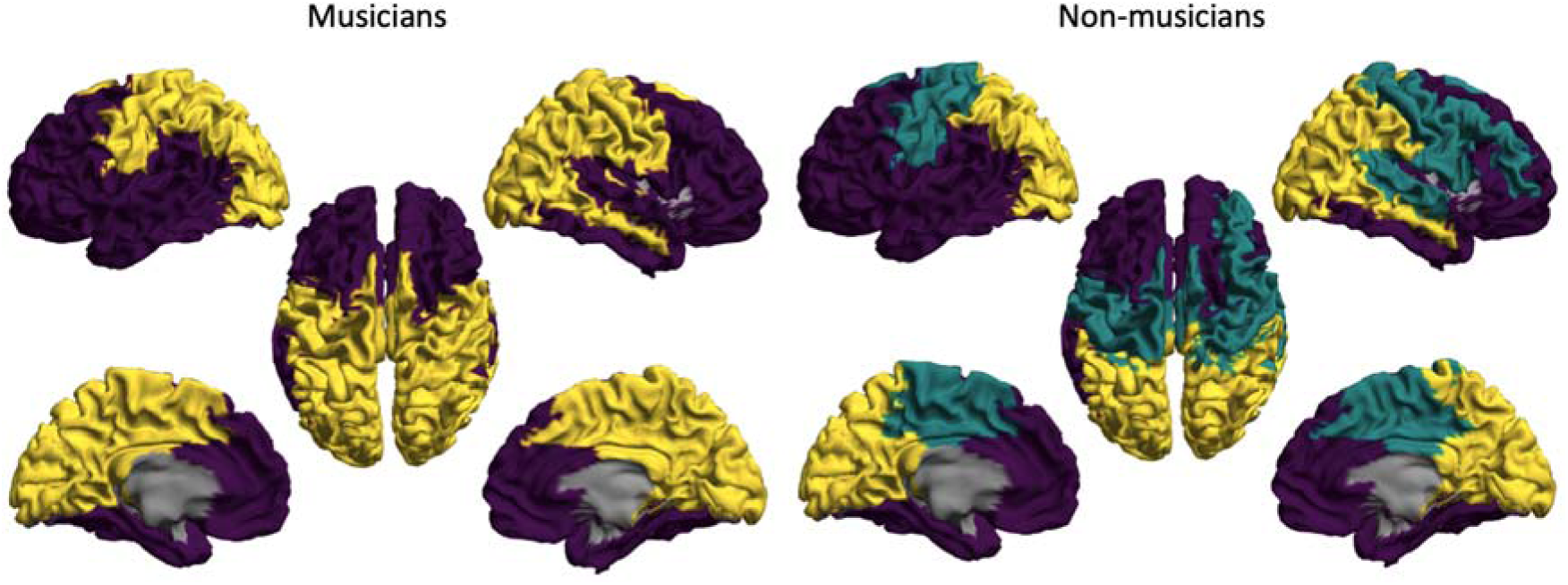
Modular structure in musicians and non-musicians. Within each group, regions shown in identical colour have been assigned to the same community (module). **(Left pane l**M) usicians have two modules (one purple, one yellow) over all values of interlayer coupling c. In both panels, the regions are coloured according to the module to which they were most frequently assigned across all interlayer coupling values (c=0:0.1:1), exceeding what would be expected by chance. **(Right panel)** non-musicians have three modules over all interlayer coupling values, one frontal module (purple), one motor module (green) and one occipital module (yellow).

We repeated the analysis without applying SVD normalisation and were able to confirm the results obtained with SVD normalisation. Specifically, the same number of modules were identified in the groups, and the significantly different modularity across all interlayer values remained consistent. The module compositions differed only in a few regions, preserving the overall interpretation of the three modules (Supplementary Figure S2).

Regarding the relation of modularity with other demographic factors, we found a significant monotonic relationship between age and modularity (Spearman’s rank correlation: r=-0.046 and p<0.001, over all interlayer coupling c and all runs of the community detection algorithm), meaning that aging has a modularity-decreasing effect. We also calculated the correlation between the age at which musicians started taking lessons and modularity and found a significant monotonic relation (Spearman rank correlation: r=-0.0327 and p<0.001). To further understand the link between musical training and neural connectivity findings, we correlated years of musical training with modularity and have a significant monotonic relationship between those two factors (Spearman’s rank correlation: r=-0.0319, p<0.001). Since we found a significant relation between age and modularity, we corrected this result for age via partial correlation calculation (Spearman’s rank correlation: r=-0.038, p<0.001).

## Discussion

We studied whether an impact of musical training would be detectable in musicians’ brain function, measured with MEG during resting state. To this end, a multilayer network framework to construct multi-frequency networks in a group of musicians and non-musicians was used. We studied the properties of these networks using graph-theoretical approaches. As per the hypothesis, a modular organisation in musicians spanning the visuomotor regions and the frontal and auditory parts of the brain was found. This modular organisation points to a stronger inter-modular connectivity, potentially underlying the integrated organisation observed in musicians. It must be noted here that neither single layer analysis using NBS nor seed-based connectivity analysis revealed significant group differences. Therefore, our results highlight the added value of the multilayer analysis.

The present methods-focused study adds to the body of knowledge about brain reorganisation in highly-trained musicians, which was so far limited to structural MRI studies for example (Schlaug et al., 1995), (Amunts et al., 1997), (Bangert & Schlaug, 2006), and task-based for example, (Alluri et al., 2017; Grahn & Rowe, 2009) as well as resting-state fMRI studies for example (Luo et al., 2012, 2014; Zamorano et al., 2017). The question whether musical training would also be reflected in electrophysiological functional networks outside of task conditions – during resting state – has only recently been addressed, to our knowledge in a single study so far: (Klein et al., 2016). They reported an increased intra- and inter-hemispheric functional connectivity in resting-state EEG data of musicians in theta (6.5-8Hz) and alpha1 (8.5-10Hz) frequency bands. However, their investigation was limited to studying frequency specific networks in isolation. This single layer approach does not detect pan-spectral functional networks.

The findings from our multilayer analysis underscore an important aspects about network organisation in musicians: At a global pan-spectral level, musicians have less segregated clusters (lower modularity), thereby allowing for an integrated network organisation over the whole multi-frequency network. For example, (classical) musicians often need to read sheet music, encode the visual information (alpha band network) and translate it into signals to be executed by the motor cortex (beta band network). The between network coordination is not limited only to visuo-motor areas, but has also been shown to span auditory, motor and other cognitive areas, which likely facilitates higher overall network integration (Jäncke, 2012; Proverbio et al., 2014). For instance, the coordination between auditory and motor areas is said to be mediated via the arcuate fasciculus (Catani et al., 2005; Catani & Mesulam, 2008). It is also plausible that this anatomical architecture underlies the auditory-motor coupling reported in musicians (Grahn & Rowe, 2009; Jäncke, 2012).

The present study compared musicians (all with years of training in a visual-auditory-motor skill) and non-musicians, thereby complementing earlier studies of changes in modular brain organisation in the course of learning over weeks. In a resting-state fMRI study of participants who learned to perform a visuo-motor task, (Bassett et al., 2015) found segregation of visual and motor modules, increasing with the number of trials. Similarly, during improvement of task performance, segregation of modules was observed by (Wang et al., 2024) and found to be correlated with learning rate. In the same study, segregation did not correlate with habit strength. Due to the relationship between habits and motor skills (Du et al., 2022), the result reported by (Wang et al., 2024) is a possible link between findings from learning-based studies and our result of integration rather than segregation in musicians after years of training.

The critical difference observed in our analysis regarding between layer organization lies primarily in the theta-beta1 and alpha1-beta1 layers. The present results observed in lower frequency bands may reflect long-range communication (Ward, 2003). Both theta and alpha band activity are involved in a wide range of cognitive functions (review: (Başar et al., 2001; Ward, 2003) – such as memory and attention, which are highly active areas while playing an instrument (Schlaug, 2015).

The relationship between behavioural specialisation and changes in the underlying neuronal activity is not very straightforward (Zatorre et al., 2012). This problem is more pronounced when investigating spontaneous neuronal oscillations (using M/EEG), as both increases and decreases in activity can reflect a different brain state (Baillet, 2017). There are several studies (Schneider et al., 2002; Shahin et al., 2003; Strait & Kraus, 2011, 2014) that have used evoked potentials to investigate musical specialisation. However, such evidence does not exist for resting-state data. Our results also show a significant correlation between multilayer network measures with years of training as a behavioural variable. Modularity in this case might not be sensitive enough to detect correlation within the musician group and can also detect between-group differences (absence vs. presence of experience), which has been confirmed in a predictive classifier (Supplementary Material 3). A possible solution to circumvent this issue would be to conduct a longitudinal electrophysiological study to establish a clearer link between behaviour and increase or decrease in the underlying neuronal activity (using a similar approach as in the structural imaging study by (Draganski et al., 2004)). In addition, the number of analysed group members in the musician group (n=31) in our study is also small and could be the reason for not identifying a clear relationship between behaviour and neural dynamics.

Brain dynamics occur across multiple spatial and temporal scales. Various neuro-imaging techniques measure different aspects of the same underlying physiological processes, but these data sets are often analysed in isolation. However, the multilayer network framework has the potential to bridge this gap by taking a more integrative approach (De Domenico, 2017; Mandke et al., 2018). For example, the framework can be used to integrate data from MEG in different frequency ranges (as in the current study), from DTI and MEG or fMRI and MEG in the same group of participants. As a future research direction, one can also integrate the structural network information as an additional network layer in the multilayer framework, as e.g. investigated in (Battiston et al., 2017, 2018; Lim et al., 2019) for one fMRI-based and an additional structural network layer for healthy controls. Recently, a framework for including - next to structural and functional layers - also a third morphological grey matter network layer has been proposed (Casas-Roma et al., 2022). Therefore, the multilayer network approach allows us to study in an individual how brain processes at different scales, from functional to structural, undergo plasticity related changes. Our study is one of the first demonstrations that an integrative approach is useful to study training related plasticity.

The present study is not without limitations. Firstly, the data used here were part of the Open MEG Archive (Niso et al., 2016), which included a number of participants (musicians, 8 out of 31 participants), who played an instrument for 5+ years but did not identify themselves as musicians. In addition, in our group of musicians, the participants did not play the same instrument (Supplementary Table 1) nor did they have similar years of experience (Mean: 13.25 years; SD: 5.04) playing an instrument. A future study could be designed to replicate these findings, with a more careful selection of participants to identify a clear relationship between resting-state network organisation and years of experience. The neurobiological interpretation of our results should be approached with caution since a demonstration of their reproducibility in a second independent large dataset has been left as future work. Secondly, the MEG recordings in our study were only 5 min in duration. The recording duration, an important aspect in capturing brain states in individual subjects (Liuzzi et al., 2017), should be carefully considered when designing future resting-state EEG or MEG studies. The OMEGA questionnaire data does not include more specific information on musical training or preference of musical style. As a result, we were unable to investigate these aspects. With more elaborate questionnaire data, one could analyse which precise factors of musical training cause the multilayer network changes. Another limitation is the use of self-report questionnaires in the OMEGA study, which are known to inherit multiple biases (Schwarz, 1999). While this method is common in the field, future studies should consider using more objective measures to quantify musical expertise such as the short/mini-Profile of Music Perception Skills (Zentner & Strauss, 2017).

The majority of the previous studies that have reported group differences in motor cortices of musicians in both structural and task-based functional data (see for example: (Palomar-García et al., 2017; Schlaug, 2015; Schlaug et al., 1995; Stewart, 2008), have recruited participants that play the same instrument professionally and have very similar levels of expertise. A critical limitation of the present study is the heterogeneity among participants in terms of years of experience (13.25 years ± 5.04) and a wide range of musical expertise (string instruments, percussions etc). This heterogeneity coupled with resting-state MEG recordings that were only 5-mins long probably make identification of seed-based differences difficult. Hence, this methods-focused study focused on identifying global changes in the network configuration. However, it is plausible in a future study to introduce a task that engages both auditory and motor systems, which would likely make the differences more apparent.

Recently, in the context of speech, (Assaneo & Poeppel, 2018) provided evidence for the presence of an intrinsic rhythm – which mediates auditory-motor communication. It is conceivable that such a fine-tuned mechanism might also exist in musicians, mediating the communication between different areas and potentially leading to the formation of sub-networks or communities. This analysis needs to be probed further by studying the relationship between task-based data and task-free data in the same group of musicians.

To conclude, we used an open access dataset (OMEGA, (Niso et al., 2016)) to investigate the effects of musical training related plasticity in resting-state MEG data. We studied the frequency specific networks using NBS and by applying a multilayer network framework to construct multi-frequency networks in musicians and non-musicians, which is a novel framework in this context. The single layer analysis did not reveal a statistically significant difference between the two groups. However, effects of musical training were detected in the modularity of the network, which spanned visuo-motor and fronto-temporal areas that are involved in execution of a musical performance. The results also indicate that musicians show an integrated network organisation in comparison to non-musicians. Presently, the study of musical training related plasticity lacks an integrative approach and therefore neglects interactions between processes of plasticity occurring at different frequency bands and/or different modalities within individuals. To overcome this limitation, we suggest the use of the multilayer network framework.

## Data Availability Statement

The data that support the findings of this study are openly available in Open MEG Archive (OMEGA) at http://doi.org/10.23686/0015896.

## Author contribution statement

KM: Conceptualisation, Methodology, Investigation, Software, Data curation, Formal analysis, Visualisation, Writing: original draft, review, editing.

PT: Conceptualisation, Methodology, Investigation, Software, Validation, Writing: review & editing.

PA: Methodology, Supervision.

MS: Conceptualisation, Resources, Methodology, Investigation, Project administration, Supervision, Writing: original draft, review, editing.

JM: Conceptualisation, Methodology, Investigation, Formal analysis, Verification, Supervision, Project administration, Software, Writing: original draft, review, editing.

