## Supplementary for "Musicians’ brains at rest: Multilayer network analysis of MEG data"

**for**

^f^ Current address: Berlin Institute of Health, Charité – Universitätsmedizin Berlin, Berlin, Germany

^g^ Current address: Department of Neurology with Experimental Neurology, Brain Simulation Section, Charité – Universitätsmedizin Berlin, corporate member of Freie Universität Berlin and Humboldt-Universität zu Berlin, Berlin, Germany

^§^ Corresponding author:

Dr Martin Schürmann

**Supplementary Table 1: Details on the self-report of instruments from N=31 musicians.**

| Subject ID | Instrument(s) |
| --- | --- |
| sub-01 | Guitar |
| sub-02 | Guitar |
| sub-03 | Guitar, Bass, Drums |
| sub-04 | Piano, Flute |
| sub-05 | Guitar |
| sub-06 | Guitar, Drums |
| sub-07 | Bass, Guitar |
| sub-08 | Guitar, Violin |
| sub- 09 | Guitar |
| sub-10 | Violin |
| sub-11 | Piano, Keyboard |
| sub-12 | Clarinet |
| sub-13 | Piano |
| sub-14 | Piano, Clarinet |
| sub-15 | Cello, Voice, Piano |
| sub-16 | Piano, Bass, Drums |
| sub-17 | Piano, Saxophone |
| sub-18 | Voice, Piano, Guitar |
| sub-19 | Piano |
| sub-20 | Trombone, Flute |
| sub-21 | Drums, Guitar |
| sub-22 | Piano |
| sub-23 | Guitar, drums |
| sub-24 | Piano |
| sub-25 | Guitar, Piano, Drums |
| sub-26 | Recorders, fiddle |
| sub-27 | Piano |
| sub-28 | Piano |
| sub-29 | Piano, Saxophone |
| sub-30 | Ukulele |
| sub-31 | Flute |

**Supplementary material 1 -**

*Methodological considerations for multilayer networks: A note on interlayer coupling*

In this work, a frequency-based multilayer network was used, where individual layers of the network were derived from individual MEG frequency bands that were identified *a priori*. The intra-layer (within layer/band) communication was quantified using AEC and the inter-layer or cross-frequency (between layer/band) communication was set to one-to-one (or parameter *c*). Presently, there is no consensus in the network science community on the appropriate way of estimating interlayer coupling parameters and it suffers from some level of arbitrariness (Tewarie *et al.*, 2021). This problem is further compounded by the fact that interlayer coupling between frequency-specific networks can be determined by many different metrics. For instance, the commonly described cross-frequency interactions as per (Jensen and Colgin, 2007) are – (a) power to power, (b) phase to phase, (c) phase to frequency, and (d) phase to power. There is mounting experimental evidence for the presence of phase to power or phase-amplitude coupling both in human (Cohen *et al.*, 2009; Onslow, Bogacz and Jones, 2011) and animal electrophysiology (Wulff *et al.*, 2009). Furthermore, (Jensen and Colgin, 2007), argue that gamma oscillations (>80 Hz) tend to couple strongly with the phase of the theta cycle (4-8 Hz), which has primarily been shown in intracranial data. A similar relationship has also been shown in resting-state MEG data (Florin and Baillet, 2015), however, the highest frequency band under consideration in the present work was 16.5-20 Hz and the lowest was 6.5-8 Hz, where such a cross-frequency relationship has not been demonstrated before, making estimation of interlayer coupling a non-trivial problem. In a multilayer description, it is possible to include interlayer link weights derived from a separate measure of connectivity, such as, phase-amplitude coupling, but this will likely lead to inconsistent results and requires a careful methodological investigation. Moreover, the non-trivial problem of interlayer coupling is accentuated when considering multilayer networks where individual layers are derived from different imaging modalities (for example, layer 1: fMRI network, layer 2: MEG network).

As problematic as the interlayer coupling is, (De Domenico, Sasai and Arenas, 2016) argue that presence of such interlayer links allows us to take full advantage of tensorial algebra, allowing us to extend single-layer network metrics to multilayer networks. They also suggest that the choice of “*c*” should therefore depend on the choice of analysis. Recent work on interlayer coupling using a network reconstruction algorithm demonstrated that interlayer coupling is indeed dominated by one-to-one layer coupling for alpha to beta band while strong widespread long-distance interlayer links should be considered when coupling theta and gamma band layers (Tewarie *et al.*, 2021).

In the end, building on the work of (De Domenico, Sasai and Arenas, 2016) and to maintain consistency between findings reported by (Guillon *et al.*, 2017; Yu *et al.*, 2017) the inter-layer coupling parameter was set to *H_kl_ = cI*. However, future work should consider exploring different interlayer couplings due to their recently discovered frequency-specific nature (Tewarie *et al.*, 2021).

**Supplementary material 2 –**

The reasons for normalising the symmetric and square block adjacency matrix (as studied here) by its *largest* eigenvalue are as follows –

1) The largest eigenvalue *λ_1_* is called the spectral radius of the graph (*λ_1_, λ_2_, λ_3_ ,..., λ_n_* is the eigenspectrum). Previous work has shown that the epidemic threshold (critical threshold) of a network is given by the inverse of the spectral radius of the network’s adjacency matrix (Kivelä *et al.*, 2014; Van Mieghem, 2016)**.** Moreover, (Van Mieghem, 2016) has demonstrated that it is also the threshold of phase transition of virus spread and synchronisation of coupled oscillators in networks. Therefore, by normalising the adjacency matrix by its spectral radius, the critical threshold is equalised between groups and datasets. The inverse of spectral radius is a measure of synchronisation threshold, it is related to shortest path, overall connectedness and degree diversity of a graph (for review see (Stam and van Straaten, 2012)).
2) The largest eigenvalue is an upper bound for the norm of the adjacency matrix (Van Mieghem, 2016)**.** Therefore, normalisation based on the first eigenvalue is a normalisation factor related to the norm of the adjacency matrix. Any combination of set of values in the eigen spectrum (i.e., sum over all eigenvalues or over more than one of the highest eigenvalues) does not have such a proven mathematical relationship with the matrix norm (to the best of our knowledge).
3) Lastly, the first eigenvalue also captures the main mode of variation of functional connectivity. (Tewarie *et al.*, 2016) demonstrated this by removing the first eigenvalue and calculating the average correlation over all the layers of the multilayer network. This difference led to a sharp increase in global correlations, potentially indicating that the largest eigenvalue is likely responsible for frequency specific information.

**Supplementary Material 3 –**

*Predictive Model for Classifying Musicians and Non-Musicians*

A predictive classifier based on multilayer network feature of modularity was constructed and confirmed that we can predict whether an individual is a musician or not with above chance level (above 54%), which increased up to 100% for increasing values of inter-layer coupling (or c) (Supplementary Figure S1). We used a logistic regression model and a cut-off of at least 50% probability for being classified as musician. If we consider the average modularity over all the values of c per individual as a classifying feature, the accuracy for prediction was 96.78% (93.44% in a 4-fold cross-validation).

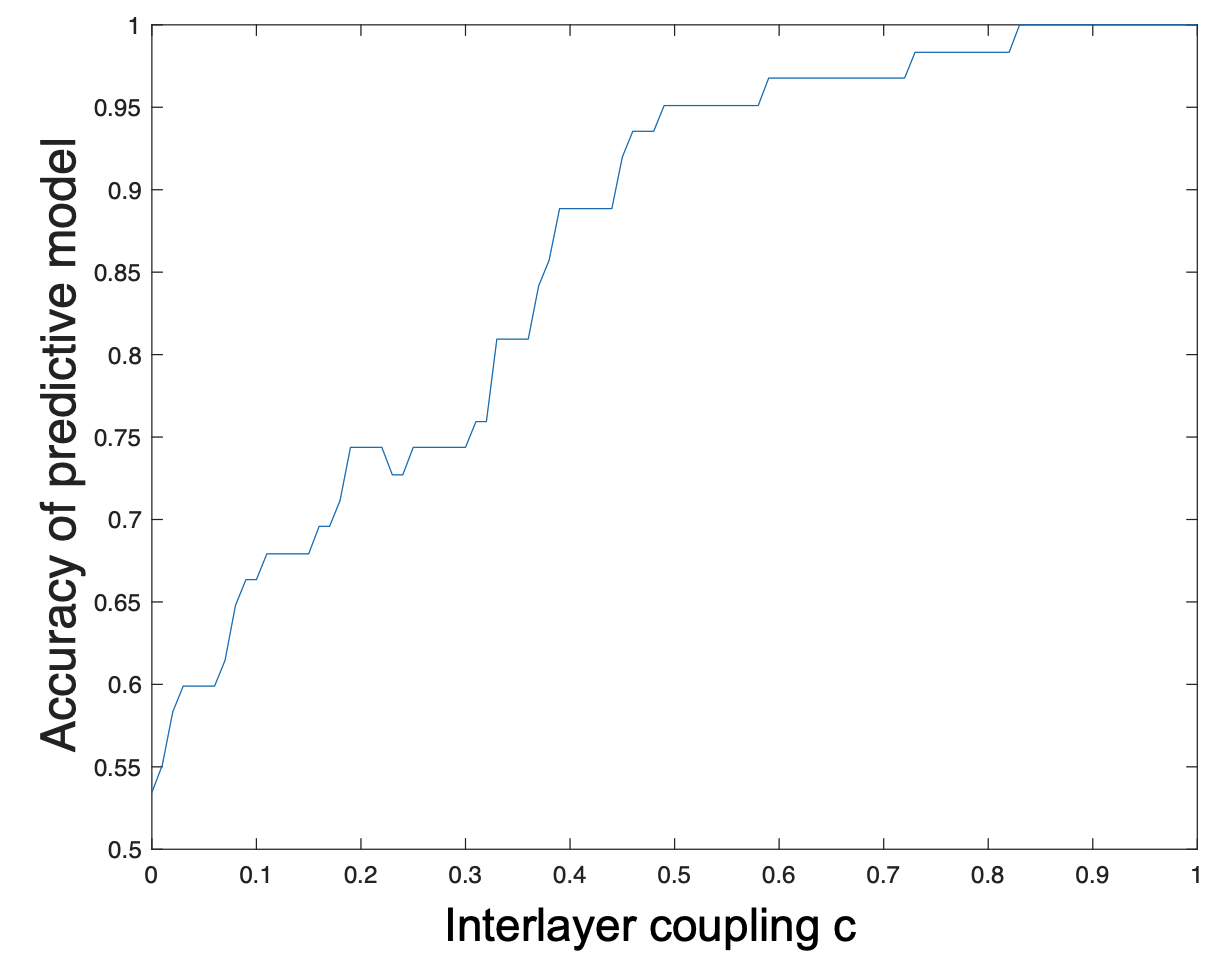

**Supplementary Figure S1: The accuracy of a predictive model over all values of interlayer coupling c.** These accuracies are based on a 4-fold cross-validation of the set of N=62 subjects.

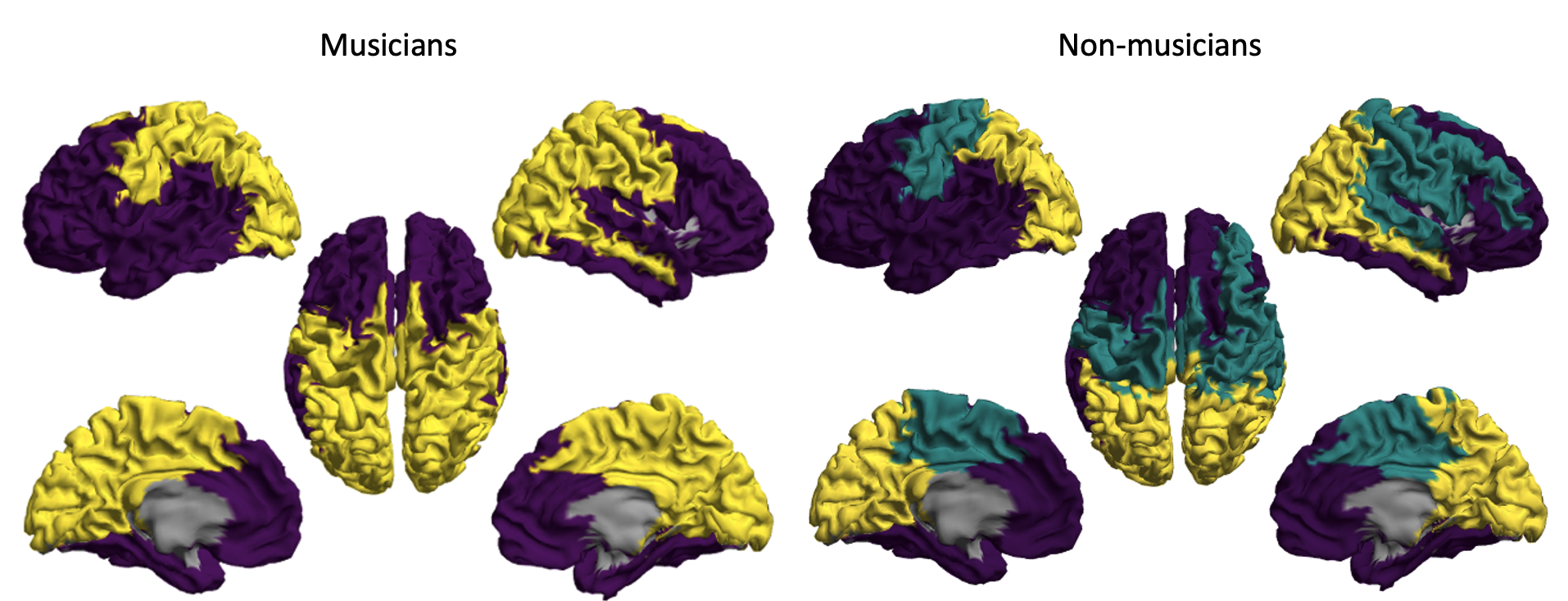

**Supplementary Figure S2:** **Modular structure in musicians and non-musicians without SVD normalisation.** Within each group, regions shown in identical colour have been assigned to the same community (module). **(Left panel)** Musicians have two modules (one purple, one yellow) over all values of interlayer coupling c. In both panels, the regions are coloured according to the module to which they were most frequently assigned across all interlayer coupling values (c=0:0.1:1), exceeding what would be expected by chance. **(Right panel)** Non-musicians have three modules over all interlayer coupling values, one frontal module (purple), one motor module (green) and one occipital module (yellow).
